# From genotype to phenotype: maintenance of a chemical polymorphism in the context of high geneflow

**DOI:** 10.1101/2020.09.24.299651

**Authors:** Bodil K. Ehlers, Perrine Gauthier, Palle Villesen, Sylvain Santoni, John D. Thompson, Thomas Bataillon

**Affiliations:** Department of Bioscience, Silkeborg, Aarhus University, Denmark; CEFE, CNRS Montpellier France; Bioinformatics Research Center, Aarhus University, Denmark; UMR AGAP INRA, Montpellier France

**Keywords:** ecologically important trait, population genomics, genetic determination

## Abstract

A major question in evolution is how to maintain many adaptive phenotypes within a species. In Mediterranean wild thyme, a staggering number of discrete chemical phenotypes (chemotypes) coexist in close geographic proximity. Plant chemotypes are defined by the dominant monoterpene produced in their essential oil. We study the genetics of six distinct chemotypes nested within two well established ecotypes. Ecotypes, and chemotypes within ecotypes, are spatially segregated, and their distribution tracks local differences in the abiotic environment. The ecotypes have undergone a rapid shift in distribution associated with current climate change. Here, combining genomic, phenotypic, and environmental data, we show how the genetics of ecotype determination can allow for such rapid evolutionary response despite high gene flow among ecotypes. Variation in three terpene-synthase loci explains almost all variation in ecotype identity, with one single locus accounting for as much as 78% of it. Phenotypic selection on ecotypes combined with low segregating genotypic redundancy and tight genetic determination leaves a clear footprint at the genomic level: alleles associated with ecotype identity track environmental variation despite extensive gene flow. Different chemotypes, nested within each ecotype, also track environmental variation. However, in contrast to ecotypes, chemotype identity is determined by more loci and show a wider range of genotypic redundancy, which dilutes the impact of phenotypic selection on alleles associated with different chemotypes. Identifying the genetics behind this polymorphism in thyme is a crucial step towards understanding the maintenance of this widespread chemical polymorphism found in many aromatic *Lamiaceae*.

## Introduction

Mediterranean aromatic herbs are with their many fragrances a striking example of intraspecific phenotypic variation (1). Most of us are acquainted with this variation: aromatic plants have been harvested by humans for centuries and we use them in pharmaceutical products, as culinary herbs and as perfume components. Thyme, Oregano, Rosemary, Lavender, and Savory are shrubs that dominate the summer-dry Mediterranean vegetation types known as garrigue, maquis or tomillares. Their aromatic essential oils are sequestered in glandular trichomes on leaves and stems. Monoterpenes are the main component of their essential oil and give the plants their characteristic scent. In many species, the identity of the dominant monoterpene in the oil varies among individuals and populations. Wild thyme (*Thymus vulgaris*) contains at least seven distinct chemical phenotypes (so-called chemotypes) (2), each chemotype producing a different dominant monoterpene, which represents 50-80% of the chemicals in the essential oil (2, 3). Many of these chemotypes are also found in other species of *Thymus* or in other *Lamiaceae* although the majority harbor fewer chemotypes^1^. Ecological roles of the monoterpenes are manifold and include deterrence of herbivores, allelopathy, and protection against abiotic stress. The chemotypes also make thyme an important community engineer: leaching of chemicals from leaves modifies the local soil environment, which affects the neighboring plant community, and these effects vary with the identity of the monoterpene (4–7).

In wild thyme, the dominant monoterpene is either a phenolic or a non-phenolic molecule. This distinction defines two ecotypes that track variation in the abiotic environment: phenolic types occur in areas with milder winters and more severe summer drought and non-phenolic thyme plants occur in areas with occasional severe early winter freezing (8). A local population is typically dominated by one or two chemotypes, and where two chemotypes coexist these are often both phenolic or non-phenolic. Long term reciprocal transplant experiments have confirmed the adaptive nature of phenolic and non-phenolic ecotypes (9). In the South of France, the spatial distribution of *T. vulgaris* chemotypes has been studied for almost 50 years (10). Chemotype variation is heritable^11^, and six distinct chemotypes co-occur within an area of 10*8 km showing a remarkable stable spatial distribution. The six chemotypes can be grouped into two ecotypes: the phenolic ecotype that consist of two chemotypes: thymol and carvacrol, and the non-phenolic ecotype that comprises four chemotypes: geraniol, alpha-terpineol, thuyanol, and linalool. Sites in close proximity (100 m) can be dominated by different eco and chemotypes. Lack of severe winterfrost in the last decades have been coupled with an increase in the phenolic ecotype and serve an example of rapid response to current climate change (10). Within each ecotype, the individual chemotypes show a remarably stable spatial distribution (1).

From an evolutionary point of view, the maintenance of several phenotypes within a species and in such close proximity is almost scandalous. How to stably maintain many discrete phenotypes within a single species is a long-standing conundrum for ecological and evolutionary theory. Theories for the maintenance of different types abound: besides trivially re-introducing new types by mutation, several mechanisms have been proposed (12). Negative frequency dependent selection, a fitness advantage to a locally rare type, automatically solves the maintenance problem. Self-incompatibility alleles in angiosperms is a known example, where numerous (mating) types are encoded by a single (super) locus; but examples of many (>2) adaptive discrete phenotypes maintained by this mechanism remain rare, but see Rock-Scissor-Paper dynamics in lizard color morphs (13). Another solution involves environmental heterogeneity, with selection maintaining locally adapted alleles conferring alternative phenotypes (14,15). Many models have been crafted to study how local selection maintains phenotypic differentiation despite drift and the swamping effect of migration (12,16,17). Here the genetics underlying phenotypic variation matters: whether one or many loci encode phenotype variation and whether these loci segregate independently or as a single entity (supergene) determines the likelihood of local selection maintaining phenotypic variation in the face of geneflow. Theory suggests that the number of genotypes that, at a given time, segregate in a population and are phenotypically redundant, so called segregating redundancy, is a critical parameter governing both the predictability of evolution (18,19) and the consequences of locally heterogenous selection for maintaining different adaptive phenotypes (20). The level of segregating (genotypic) redundancy determines the intensity of selection on individual alleles. Higher redundancy (several different genotypes can produce the same phenotype) makes it more difficult to either fix or eliminate individual alleles compared to situations where alternative phenotypes are determined by one or a few genotypes. Redundancy may also help maintain phenotypic differentiation in the face of high gene-flow, as several genotypes producing the same adapted phenotype can contribute to withstand swamping due to the immigration of divergent alleles (20).

Here, we study the genetics of chemotype variation in *T. vulgaris*. We explore how genetic determination of chemotypes and environmental variation contribute to maintain this striking polymorphism. Previous crossing experiments demonstrated that chemotype variation is strongly heritable and likely controlled by the segregation of genetic variants at relatively few (5–6) independent loci (11). More recent studies point to an effect of gene dosage with quantitative effects within chemotypes (3). However, this phenotype has never been studied using a genomic approach, and genes underlying this chemical polymorphism have not been identified.

To understand the genetic determination and maintenance of this polymorphism, we combined population genomics data with phenotypic and environmental data from 21 populations located at geographically close sites. Together, these naturally occurring populations harbor six distinct chemotypes that form two main ecotypes (supplementary Figure 1). Having so much phenotypic variation present in geographically close sites is a unique situation: It allows to study environmental selection on many different types without the confounding effect of population genetic structure that often arise at larger spatial scales. Chemotypes were determined by analyzing the chemical composition of 702 individuals (12–40 individuals per population), and of these, 252 individuals (12 from each population) were also genotyped. Environmental data were collected at each site to characterize variation in summer temperature, winter temperature, and soil properties. We use this data to study the genetic determination of eco and chemotypes, and to quantify how ecotypes, chemotypes and the alleles determining their identity co-vary with environmental variation.

## Results

### Ecotypes can be explained by genetic variation at three candidate loci and is spatially differentiated despite high gene-flow

Our genomic data consists of a set of highly reproducible genetic variants (supplementary Table 1). These are 3920 single nucleotide polymorphisms (SNPs), with a frequency of the rarest allele > 0.1. These polymorphisms are located in candidate genes of the monoterpene biosynthesis pathway and a larger set of anonymous genes (> 300 genes in total, see methods). Using a logistic regression framework accounting for the effects of underlying weak population structure (supplementary Figure 2) via five principal components describing SNP variation, and a conservative approach to account for multiple testing, we show that the two ecotypes (phenolic vs. non-phenolic) strongly associate with variation at three genetic loci. Genetic variants of one major gene - homologous to a linalool synthase - explains 78% of the observed ecotypic variation between individuals. Variation within two additional genes (homologous to two gamma-terpinene synthases) is also strongly associated with ecotypes (both p < 10^-8^), but with more modest effects (Figure 1A, Table 1A). A model combining these 3 variants accounts for virtually all ecotype variation (Figure 1A). Curiously, for two of these loci, none of the individuals are homozygote genotypes for the rarest allele (supplementary Table 2). Independent sequencing and genotyping of 23 individuals confirms that this is not an artefact of poor genotyping at these SNPs (methods, supplementary Table 1).

**Figure 1.**
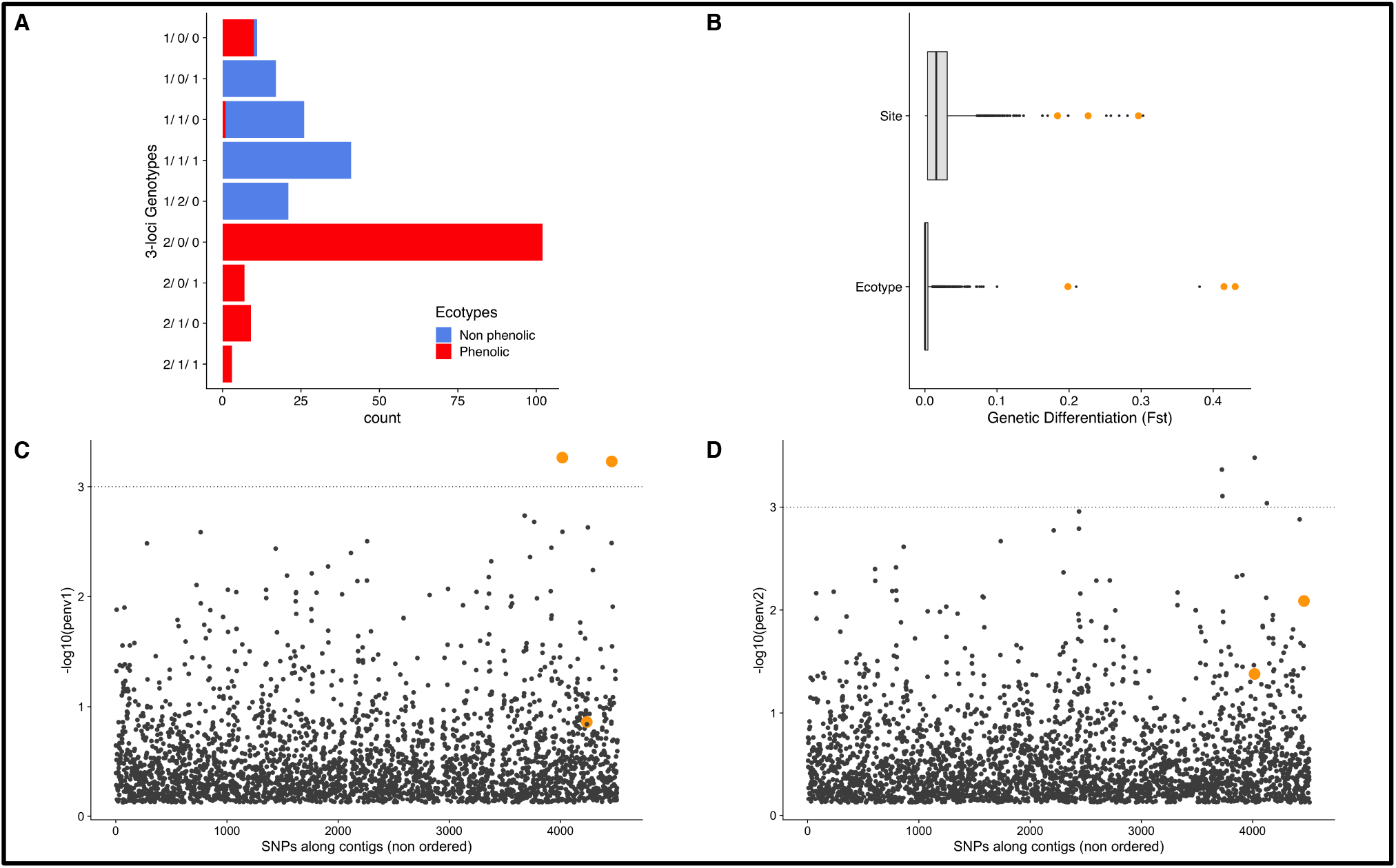
**1A**: Genotype to ecotype map. 3 loci genotypes are built using the three SNPs (contig 71000, Contig 12377, and KR920616.1) with the strongest associations (Table 1A). Count refers to the number of individuals in each category. 0, 1 and 2 denote the 3 possible genotypes at each SNP (0 and 2 are genotypes homozygous for the reference and alternative allele, 1 is the heterozygous genotype). **1B:** Boxplot of the distribution of Fst (n=3920 SNPs) among sites (n=21 populations) and between the two ecotypes. Orange dots denote the three SNPs used to build the genotype to ecotype map. **1C** and **1D**: Manhattan plots for the SNP-EnvPC1 (1C) and SNP-EnvPC2 (1D) associations. Orange dots denote the three SNPs used to build the genotype to ecotype map. For graphical convenience only p-values <0.5 are depicted (so on Fig 1D one of the orange dots is not displayed).

**Table 1.**
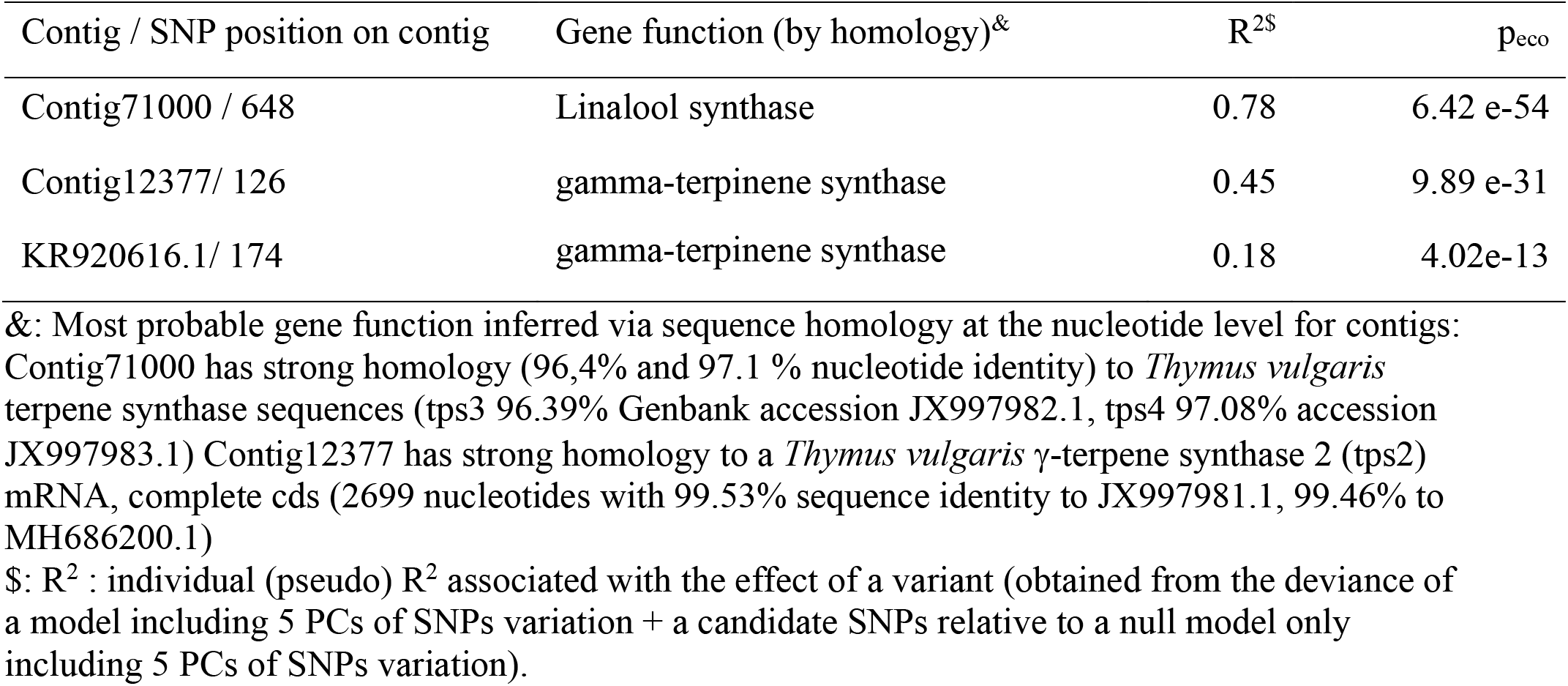
Summary of top SNP-ecotype and SNP-chemotype association. **Table 1A**: Top 3 SNP – ecotype association

**Table 1B.** Top 6 SNPs-chemotype associations. Numbers in each chemotype column refer to the p-value for a test of SNP-chemotype identity associations.

| Gene /<br>SNP position on Gene | Putative<br>Gene function | Individual Chemotype |  |  |  |  |  |
| --- | --- | --- | --- | --- | --- | --- | --- |
|  |  | C | T | G | A | U | L |
| Contig71000 /648 | Linalool synthase | 2.68e-09 | 3.06e-24 | 8.59e-05 | 4.95e-06 | 5.13e-08 | 7.9e-23 |
| Contig12377 /<br>126 | gamma-terpinene<br>synthase | 0.000264 | 1.59e-19 | 0.205 | 0.246 | 0.125 | 1.52e-30 |
| KR920616.1/ 174 | gamma-terpinene<br>synthase | 0.0017 | 8.04e-08 | 5.37e-10 | 2.62e-10 | 6.44e-14 | 1.6e-05 |
| JX946358.1 /1452 | sabinene hydrate<br>synthase | 0.0195 | 0.00845 | 0.181 | 0.25 | 1.8e-12 | 0.151 |
| KC461937.1 /1279 | alpha terpineol<br>synthase | 0.454 | 0.542 | 0.045 | 1.22e-07 | 0.00998 | 0.869 |
| KM272332.1 /359 | limonene-3-hydroxylase | 2.36e-27 | 5.01e-13 | 0.0382 | 0.968 | 0.269 | 0.00065 |

The genetic differentiation among sites, measured by average genome differentiation at the 3920 common SNPs, is very low (Fst between sites = 0.021 +− 0.003), and is even lower between ecotypes (Fst between ecotypes: 0.004 +− 0.0002, Fig 1B, C). The three SNPs displaying the strongest statistical association with ecotype variation are also among the SNPs exhibiting the highest genetic differentiation among sites (Fig 1B), consistent with strong local selection on ecotypes.

Testing the importance of both drift and environmental variation on ecotype distribution, we find the largest effect of environmental variation (Table 2).

A few genotypes account for ecotype variation and redundancy is low with respectively 2.1 (+− 0.03) and 2.4 (+− 0.06) effectively segregating genotypes determining the non-phenolic and phenolic ecotype (supplementary Table 3). These findings together with the tight genetic determination mean that strong phenotypic selection translates into strong selection on individual loci. This explains how differentiation of ecotypes between abiotic environments can be maintained despite considerable gene-flow.

**Table 2.** Association of ecotypes and chemotypes with environment after accounting for genetic drift.

| Models Fitted<sup>s</sup> | All Chemotypes | Ecoty<br>pes. | G | aT | U | L | C | T |
| --- | --- | --- | --- | --- | --- | --- | --- | --- |
| Dev(Null) | 853.30 | 103.6<br>6 | 62.36 | 52.64 | 69.01 | 85.00 | 102.64 | 104.36 |
| Dev(Genetic drift) | 737.16 | 95.71 | 35.78 | 46.16 | 54.24 | 65.95 | 76.39 | 98.07 |
| Dev(Genetic Drift + Environment) | 569.84 | 79.91 | 34.19 | 28.07 | 28.72 | 56.19 | 51.30 | 83.31 |
| R<sup>2</sup> genetic drift | 0.14 | 0.08 | 0.43 | 0.12 | 0.21 | 0.22 | 0.26 | 0.06 |
| R<sup>2</sup> genetic drift + environment | 0.33 | 0.23 | 0.45 | 0.47 | 0.58 | 0.34 | 0.50 | 0.20 |

**Models fitted:** Genetic drift model uses 5PCs of SNP variation, the Genetic drift + environment model uses 5 PCs of SNPs and 3 PCs of environmental variation to predict the distribution of all chemotypes jointly (multinomial logistic regression), individual ecotype or chemotype (G A U L C T). **R<sup>2</sup> :** R<sup>2</sup> are pseudo R<sup>2</sup> quantifying the percent of variation explained by genetic drift (R<sup>2</sup> drift) and genetic drift+ environment. These are calculated as 1 – Dev\_Drift/Dev\_Null, % explained by environment calculated as 1 – Dev\_Drift+Env/ Dev\_Null, where Dev is the deviance of each model.

### Genetic determination and differentiation of chemotypes

In the same way as ecotypes, chemotypes show very low overall genetic differentiation (mean Fst among chemotypes = 0.007 +− 0.0003, Figure 2B). We tested for associations between genetic variants (SNPs) and individual chemotype identity. When using the same conservative threshold for significance of associations (requesting p < 0.05/3920), we detected again the three variants already associated with ecotype differences, as well as additional SNPs located in three independent genes that strongly associate with individual chemotypes. The SNPs displaying the strongest associations with chemotypes are located within six genes all identified as homologs for previously reported enzymes of the monoterpene biosynthetic pathway (Table 1B). Collectively 45% of the variation in chemotype identity can be explained by variation in genotype at these six loci (Figure 2A). In addition, the number of effectively segregating genotypes underlying each chemotype vary considerably (Figure 2A, supplementary Table 3) compared to ecotypes.

**Figure 2.**
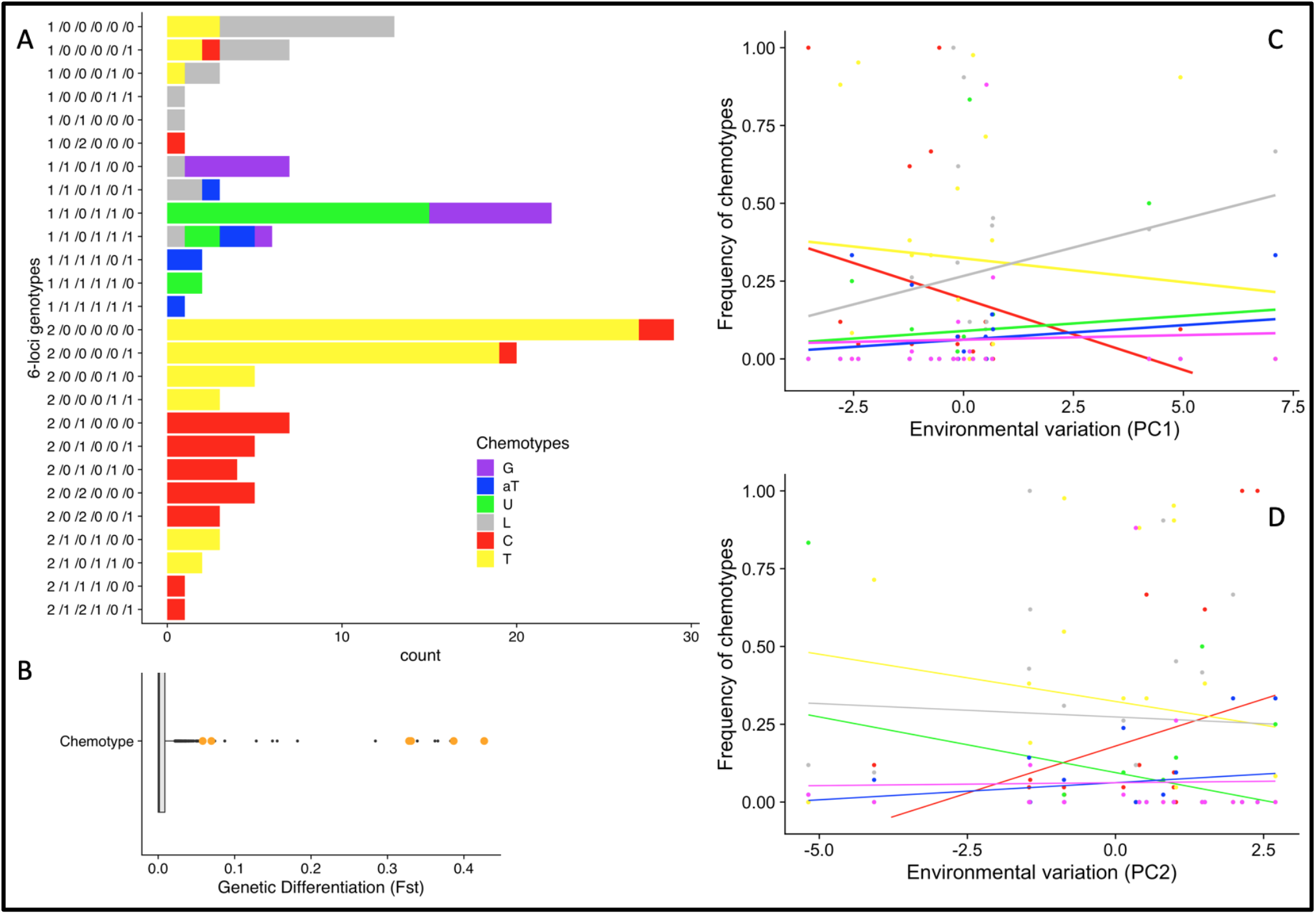
**2A**: Genotype to chemotype map. Six-loci genotypes are built using the 6 SNPs with the strongest associations (Table 1B). Counts refers to the number of individuals in each genotype category. Colors indicate chemotype identity. **2B:** Boxplot of the distribution of Fst (n = 3920 SNPs) among chemotypes. Orange dots denote the six SNPs used to build the genotype to chemotype map. **2C and 2D:** Chemotype environment associations. Each dot represents the observed frequency of chemotype, lines indicate regression lines from logistic regression, and colour of line indicate chemotype identity

The distribution of individual chemotypes across populations is by far explained by environmental variation (principal components), with a comparatively smaller effect of genetic drift (Table 2). Individual chemotypes within ecotypes are not interchangeable and track differently the environmental variation (Figure 2, Table 2). The six SNPs that have the strongest association with chemotypes were also among the SNPs that showed most differentiation among sites (Figure 2B), suggesting that although gene-flow among populations is high, heterogeneous selection caused by variation in local abiotic environment also drives the spatial segregation of chemotypes within ecotypes. One notable exception here is the “geraniol” chemotype (G) that is the rarest chemotype in the study area but locally very frequent in two sites. For this chemotype, the variables measuring local drift (principal components of SNPs variation) and environmental variation are confounded (Table 2).

### Effects of selection on genetic variation depend on the genetic architecture of phenotypes and their redundancy

If ecotypes, and chemotypes nested within ecotypes, track environmental variation, we expect genetic variants underlying this phenotypic variation to be associated with environmental variation. We examined the extent to which SNPs were associated with environmental variation. We characterize the environment using PC1 and PC2 of environmental variation that account respectively for 45% and 28% of the total environmental variation (supplementary Figure 3). Using a logistic regression framework that also accounts for local genetic differences (using 5 PCs of SNPs variation as above), we found only a few SNPs with suggestive SNP-environment associations (Figure 1 C-D) but none of these are statistically significant after correction for multiple testing across all common SNPs. Among the six major SNPs determining eco- and chemotype identity (Table 1, Figure 1A and 2A), the three SNPs that explain ecotype identity, are also those with the strongest changes in allele frequency across environment (supplementary Figure 4, supplementary Table 4). These SNPs (contig 71000, and especially Contig 12377, and KR920616.1, Table 1A) show association with the abiotic environmental variation captured by PC1 of environmental variation. This is expected given that the two phenolic types, Carvacrol and to a lesser extent Thymol, covary most with PC1: phenolic types are rare at sites with harsh winters (Figure 2C).

We note that our sampling design relies on environmental variation at 21 sites, which lacks statistical power to detect weaker SNP-environment associations. It may be argued that some SNP - chemotype associations could be caused by population structure not accounted for by the five SNP PCs. However, these associations still hold even when accounting for population structure by using up to ten SNP PCs (supplementary Figure 5). Moreover, if such associations were a mere artefact of population structure, we would not expect SNPs associated to ecotypes and chemotypes to be always located in genes of the monoterpene pathway, as found in our study.

## Discussion

We demonstrate how ecotypic differentiation can be maintained in the face of considerable gene flow. Ecotype identity is under oligogenic control and variation at a single gene explains the majority of ecotype variation. This has important consequences for how selection operates on phenotypic variation to maintain adaptive differentiation. The loci accounting for most of the heritable variation in a fitness related trait are expected to undergo the strongest selection at the genotype level (21, 22). Accordingly, allele frequencies of SNPs located in the three genes for ecotype differentiation also follow the environmental variation (env PC1) that is tracked by ecotypes. The finding of major effect genes, combined with low genotype to phenotype segregating redundancy, can explain the rapid increase in the frequency of the phenolic ecotype in relation to climate warming recently documented in the same study region (10).

Unlike ecotypes, the frequency of SNPs displaying strong associations with chemotype identity had much weaker associations with environmental variables. Theory predicts that the effect of selection on a locus controlling phenotypic variation for a fitness related trait can be influenced by selection jointly acting on other loci (21–23), and that redundancy of the genotype to phenotypes has a profound impact on the evolution and maintenance of polymorphisms (20). The amount of segregating genotypic redundancy controls how phenotypic selection will leave (or not) detectable footprints at the genome level. In our study system, ecotypes are determined by few loci and a low segregating genotypic redundancy (Figure 1A), whereas chemotype determination involves more loci and more variable segregating genotypic redundancy (Figure 2A). The consequence is that genetic variation at the loci contributing to ecotypic variation more closely aligns with environmental variation, whereas this signal is much weaker at the loci that account for a comparatively smaller fraction of chemotype variation. In addition to affecting how phenotypic selection may leave a detectable footprint at the genomic level, genotypic redundancy can also provide robustness to swamping by migrating alleles. This may contribute to maintain the high number of chemotypes despite the high gene flow observed throughout our study region.

Unravelling the genetics of wild thyme chemical polymorphism is a crucial step towards understanding the evolution and maintenance of this ecological important polymorphism also found in many other sister species and genera in the *Lamiaceae*. It open the possibility to test if variation at the same genes controls chemotype variation in other aromatic plant species. This allows testing if chemotypes arose independently or by reusing genetic variation that may predate speciation.

## Materials & Methods

### Plant material

In June 2016, 21 sites were sampled within the Saint Martin de Londres area (supplementary Figure 1). The sampling of sites was based on prior knowledge of the chemotype variation in the area. Sites were chosen to maximize the diversity of abiotic environment as well as to span the known spatial gradient of chemotype diversity in the region. At each location, 12 individuals were sampled, ensuring that individual were growing at last 1,5 m apart. For each individual two types of sampling were done: fresh leaves were gathered in sealed plastic bags and stored in cooling bags until returned to the labs where samples were kept at −80 degree C until DNA extraction. In addition, leaves (shoot tip app 2 cm) for GC-MS analysis were sampled in 1 ml of methanol. After 24 h, the methanol extractions were transferred to fresh tubes and stored at −80 degree C until analyzed on GC-MS (10).

### Phenotyping: Chemotypes

The chemotype of the plants sampled from the 21 sites (252 individuals) was determined as the identity of their dominant monoterpene (either geraniol, alpha-terpineol, thuyanol, linalool, carvacrol, or thymol) determined from GC-MS. In brief, methanol extracted samples were analyzed on a ShimadzuGCMS-QP21010 fitted with a flame ionization detector and a fused silica capillary column SLB-MS (30 × 0.25 mm; 0.25-μm thickness). Helium was used as a carrier gas. The dominant compounds were identified by comparing retention time and mass spectra with standards from Mass spectral library. See Thompson et al (10) for more details.

To increase information on chemotype frequency within and among sites, we added available data on individual plants sampled from the same sites that were chemotyped just few years prior to our sampling. Our data set representing chemotype frequency at the 21 sites thus consists of chemotype identity of 702 individual thyme plants (252 individual sampled in 2016, and 450 individual plants from Thompson et al (10).

### Environment variables

#### Soil variables

At each site, 1 kg of soil was collected with 2-3 different representative samplings within each site. Soils were dried at 40°C for 48 h, sieved at 2 mm and stored in a cool room prior to analysis. Water retention potential (WRP) was calculated as the percentage of water lost after drying a wet soil for 48 h at 40°C. Water retention capacity (WRC) was then calculated as the percentage of water remaining in this previously 40°C-dried soil by a repeated drying of the sample at 110°C for 5 h. Organic matter (OM) was estimated as the percentage of matter lost after burning a dried sample at 500°C during 5 h.

#### Temperature variables

At each site, two data loggers were placed from Jan-Feb, and from May-September. Data loggers were secured on branches of thyme or other small shrubs ca. 20-40 cm above the ground. Data loggers recorded temperature every 30 min. A few data loggers were lost. After collecting data loggers, the recorded temperature was extracted and, for each site, used to create the temperature variables. The individual variables used were percentage water in soil, soil water retention and percentage organic matter in the soil (“pctsoilwater” “pctwaterret” “pctorgmat”), the mean daily minimum and maximum temperature in winter (“mint” “maxt”), the minimum temperature in the coldest month (“mintjan”), the number of days where moderate to strong freezing was recorded (below −8C) “freezemoderate”, the mean daily minimum and maximum temperature in the summer (“maxsumt”, “meansumt”), the number of days exceeding 40C or 45C (“dayabove45”, “dayabove40”), and the mean summer daily temperature amplitude (“meansumamplit”). For summer temperature, we omitted the data recorded between 12:00 and 16:00, as this represent a time of the day where direct sunlight may have caused data loggers to record unusually high temperatures not representative of the site. We checked that omitting these time intervals did not alter the ranking of the warmest to coldest sites.

### Locus targeted genotyping

We used a locus targeted genotyping procedure. A set of 20 000 DNA baits (myBaits, Arbor Biosciences) were designed to target a set of candidate genes identified by transcriptome sequencing (24) as well as candidate for known genes encoding enzymes of the monoterpene biosynthesis pathway previously identified in *T vulgaris* or close relatives (Genebank Accessions: JF940523.1, KM272332.1, KM272331.1, KU699534.1, KU699532.1, KU699531.1, KU699530.1, KU699529.1, KR920616.1, KC461937.1, JX946358.1, JX946357.1).

Plant DNA was extracted from 15 mg of fresh young leaves. Genomic library preparation for multiplexed individuals and enrichment step by bait capture follow published protocols (25, 26) with some modifications (see details on the protocols used are available as supporting information text). Briefly, for each genotype a genomic barcoded library was built. 48 barcoded libraries were mixed together for the capture by hybridization step. To increase the specificity of the enrichment step, we used a capture, washing and re-capture procedure. Captured DNA was sequenced by sets of 144 genotypes on two runs of HiSeq 3000 Illumina NGS sequencer.

### Read alignment and SNP calling

All reads were individually mapped with BWA (27) against a reference sequence compiled from two sets of sequences. The first set was a full de novo transcriptome assembly (24). The second set was Genbank reference sequences from a list of candidate genes (see list of accessions above). Each final sequence in the reference were tagged as either “full candidate” (FC: and the Genbank sequences), “anonymous target” (AT) or “background” (BG). All mapped reads were used and SNPs called using read2snps v. 2.0 (28) and GATK v. 3.8 (29) using default settings.

### SNP quality control and SNPs genotyping

We only used SNPs that were detected by both SNP calling methods (GATK and read2snps) and we also restricted our attention to SNPs with a minor allele frequency equal or greater to 0.1 (averaged over all individuals). This yielded 4520 commons SNPs. The choice of minor allele frequency threshold was to ensure both reliability of SNPs genotypes but also to have minimum power for detecting association between a SNP genotype and chemotype (or environmental) variation.

### Repeatability of SNP genotypes

We used twenty-three technical replicates (i.e. individual plants with the same DNA extraction but completely independent myBAITS enrichment, and library preparation, sequencing, read mapping and variant calling) to assess the overall reliability of sequencing and SNP calling.

We first computed the proportion of the time the exact same genotype was called (perfect genotype recall) across the technical replicates. We found that common SNPs (minor allele frequency of 0.1 or higher) were highly reproducible (mean proportion of perfect recalls: 0.996, median of 1). As a complementary check, we computed the correlation of scores on the first 10 PCA coordinates for the replicates. Doing so confirms the overall reproducibility of our sequencing, SNP calling and SNP genotyping on the set of common SNPs (Supplementary Table 1). Last, we checked that the proportion of perfect genotype recalls for the 6 SNPs displaying the highest SNP-chemotype associations (Table 1B) and also confirmed a high reproducibility for these SNPs (109 perfect genotype recalls out of 110 genotypes called).

### Principal components of SNPs variation and genetic differentiation

In order to summarize the patterns of genetic variation between common SNPS, we performed a principal component analysis (PCA) on the genotypes scores of all common SNPs. Each individual SNP genotypes were centered and reduced (mean zero and unit variance) before performing a PCA. A probabilistic PCA, as implemented in the R package *pcaMethods* (30) was used and 10 PCs were computed. The PCs were used for accounting for population structure when detecting SNP-phenotype (ecotype or chemotype) and SNP-environment associations (see below).

To quantify genetic differentiation between population, ecotype and chemotypes, we estimated Fst (using Weir and Cockerham estimators) as implemented in the R package NAM (31) using a set of non-redundant common SNPs. A set of non-redundant 3920 SNPs was obtained using the quality control function provided by the package NAM (requesting no more than 50% missing data per SNP, and excluding highly correlated SNPs, threshold 0.98).

### SNPs-Ecotype and SNPs-Chemotype association

We tested for associations between common SNPs and chemotype (or ecotype) identity across the sample of individuals of the 21 sites. We used a logistic regression framework where we are evaluating association between a SNP genotype and a binary phenotype (here a given chemotype or ecotype identity) by comparing the fit of two models:

A background model (hereafter M0) where 5 PCs of common SNPs variation are used to predict chemotype:

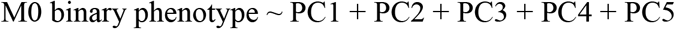

We expect M0 to have some predictive power because of the genetic differentiation between populations and the fact that populations differ on average for their ecotype (chemotype) composition. A SNP model (hereafter M1) that also uses a single SNP genotype (coded as 0,1,2) as predictor

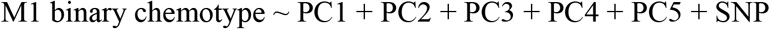

M1 models were fitted by maximum likelihood for each common SNP. This was done using the *glm*() function in R and model M1 was implemented using a logistic link and a binomial distribution for the binary phenotype.

The phenotypic effect associated with an individual SNP was computed using the (pseudo) R^2 associated with model M1 for this SNP. As M0 and M1 are generalized linear models, several ways to compute R^2 have been proposed and there is no agreement on an obvious “optimal” R^2 measure (32). We computed R^2 as 1 - Dev(M1)/Dev(M0), where Dev(Mi) is the deviance of model Mi. In brief the rationale for this choice is as follows. Deviance of M0 is analogous to a residual sum of square in a linear regression model: the bigger the lack of fit the bigger Dev(M0). Accordingly, deviance of M1 measures the lack of fit of model M1. This measures the improvement in our ability to fit individual (binary) phenotypic values across individual when we use the extra genotype information brought by a single SNP.

To test for association at each SNP and obtain a formal p-value for association, we used a likelihood ratio test and used twice the difference in deviance between M0 and M1 (Gobs = 2 (ln M1 – ln M0)) as test statistic. We assumed that under the assumption that data comes from model M0, we expect Gobs to be chi-squared distributed with one degree of freedom. Using the background model M0 as null model guards against spurious SNP-chemotype association merely due to the fact that there is some (weak) genetic differentiation between populations and differences in chemotype/ecotype frequencies across populations.

Just as in the calculation of genetic differentiation (Fst), we used 3920 SNPs, where the minor allele frequency was above 0.1, and all individuals that were both genotyped and phenotyped (n=248 after quality control on SNPs genotyping) to detect associations.

We used a strict Bonferroni correction to account for the fact that associations between SNP and the binary ecotype and chemotype were tested at numerous SNPs. Here, given that nT= 3920 SNPs were tested, α/nT is used as significance cutoff for each individual SNP. Setting α = 0.05, this amounts to require an individual SNPs significance threshold 0.05/nT= 1.27551e-05.

Visuals checks (not shown) were performed on the empirical distribution of p-values for each phenotype to ensure that these were properly calibrated. We expect most p-values to conform to M0 and accordingly a uniform distribution in [0,1] for p-values with a minor bulge of lower p-values (for SNPs displaying associations as specified underM1).

Finally, we checked the robustness of our approach by recomputing SNP-ecotype associations using 10 PCs of SNPs variation (instead of 5) in implementing M0 and M1. The rationale for doing so is to validate (*a posteriori*) that our modelling approach is not sensitive to the choice of the number of PCs included in M0 to account for population structure.

We checked that SNPs scores (defined as −log10(p-values)) computed either method were highly correlated (observed correlation 0.98, p-value < 2.2 10^-16^). We also performed a visual check to ensure that top p-value ranking was insensitive to the modelling choice of 5 vs 10 PCs (supplementary Figure 5).

### Segregating redundancy of genotypes controlling phenotypes

A set of *T* multilocus SNPs genotypes *G_i_*, i = 1..*T*, with genotypic frequency *p_i_* segregate 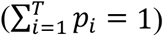, and can produce a set of discrete phenotypes. More precisely, *K* (heritable) discrete phenotype *P_j_* (*j* in {1..*K*}) are “encoded” by the set of genotypes *G*_1_,..*G_T_*. We define a coefficient *w_ij_* that, for each pair (*G_i_, P_j_*), measures how strictly phenotype *P_j_* is determined by genotype *G_j_*. In other words, the encoding might be genetically strict or loose depending on the heritability of phenotypes. We use the conditional probability that genotype *G_i_* will produce phenotype *P_j_* as a way to quantify how strict is the genotype to phenotype encoding: *w_ij_* = *Prob*(*P_j_\G_i_*). Note that *w_ij_* = 1 captures the case phenotype *P_j_* is strictly genetically determined by a specific genotype *G_j_*. Inspired by a recent review of measures of redundancy (20), we propose a measure of segregating redundancy for each discrete phenotype *P_j_* based on the effective number of genotypes, akin to Nei’s effective number of alleles via Nei’s diversity, that can produce it. We also introduce our weighting *w_ij_* when calculating this effective number of genotypes. We first calculate a weighted genotypic diversity that is segregating for determining phenotype *P_j_* as: 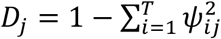, where 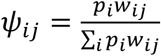

The segregating redundancy underlying Phenotype *P_j_*, *SR_j_*, is defined as the effective number of genotypes encoding *P_j_*. Accordingly, we compute *SR_j_* as the reciprocal of the weighted diversity 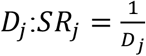. If many genotypes capable of producing *P_j_* exist in a population, *SR_j_* is maximal when all genotypes are equally frequent and encode the phenotype *P_j_* with identical probability. If only a few genotypes *G_i_* encode *P_j_* with high probability, then *SR_j_* will be much lower. We assume *n* individuals are genotyped and phenotyped (for chemotype or ecotype identity), we used the sample frequency of genotypes and phenotypes to estimate *p_i_*’s and the *w_ij_* and, accordingly, *ψ_ij_* and *SR_j_*’s. Resampling by boostrap at the individual level is used to obtain the standard errors around *SR_j_* estimates.

### Ecotype/chemotype - Environment association

The chemical phenotypes of 702 individual plants were used. To characterize environment variation, we first performed a principal component analysis on the environmental variables recorded at each of the 21 sites of our study. All environmental variables were centered and reduced before PCA. PC1, 2 and 3 account respectively for 45%, 28% and 12% of the total variation measured across sites (supplementary Figure 3).

We used a (multinomial) logistic regression, where counts of individuals in each ecotype (phenolic C,T versus non-phenolic GAUL) or all six chemotypes (GAULCT) at each site are used as response variables and the principal components of environment variation are used as predictors. Likelihood ratio tests comparing the deviance of models were used to obtain statistical significance level for the effect of the different (principal) components of environment variation on ecotype distribution. Likelihood ratio tests were conducted assuming that, under the null hypothesis of the reduced model being correct, differences in likelihoods are accurately approximated by a chi-squared distribution (with p degrees of freedom if models differ for p fitted parameters). To quantify the amount of variation accounted by each model Mi, we also used the pseudo R^2 of model M_i_ computed as 1 – Dev(M_i_)/Dev(M_0_) where Dev(M_i_) is the deviance of model Mi and Dev(M_0_) is the deviance of a null model with no environmental variable as predictors and merely fitting an intercept (equivalent to assuming a constant proportion of phenolic ecotypes across all sites).

### SNPs-environment association

To detect SNP-environmental association, we used a similar logistic regression framework as the one described above for SNP-phenotype associations. Here, instead of using phenotypes as response variables, we model variation of SNP frequency from site to site (our response variable) and test whether SNPs covary with the local environment.

Similar to above, a background model (M0) where 5 PCs of common SNPs variation is used to predict allele frequency variation at a focal SNP:

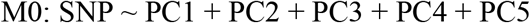

An environmental gradient model that also uses an environmental covariate (coded as a continuous variable) as supplementary predictor of the SNP frequency.

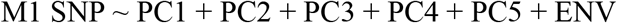

A likelihood ratio comparing M0 and M1 test is again used to test for SNP-ENV association and obtain a p value significance of the association. Using the background model M0 with 5 PCs as null model guards against spurious SNP-ENV association due to genetic differentiation between sites and differences of local environment across sites. We fitted (M0, M1) models using each common SNP as response variable and using separately PC1 and PC2 of environmental variation as ENV covariate.

All generalized models described above were using multiple predicting variables (such as PC of SNPs and PCs of environmental variation as well as individual SNPs genotypes) and were checked for correlation between predictors by computing variance inflation factor (32) implemented via the *car::vif()* function R package *car* version 3.0-8 (33).

## Supporting information

Supplementary information (incl. tables figures and methods)

## Acknowledgements

Authors thank R. Kassen for his comments on the manuscript, and T. Lenormand, M. Schierup, and R. Kassen for helpful discussions about this work.

## Notes

### Competing Interest Statement

The authors have declared no competing interest.

