## Supplementary information (incl. tables figures and methods) for "From genotype to phenotype: maintenance of a chemical polymorphism in the context of high geneflow"

**This PDF file includes:**

- Figures S1 to S5
- Tables S1 to S3
- Supplementary text
- SI References

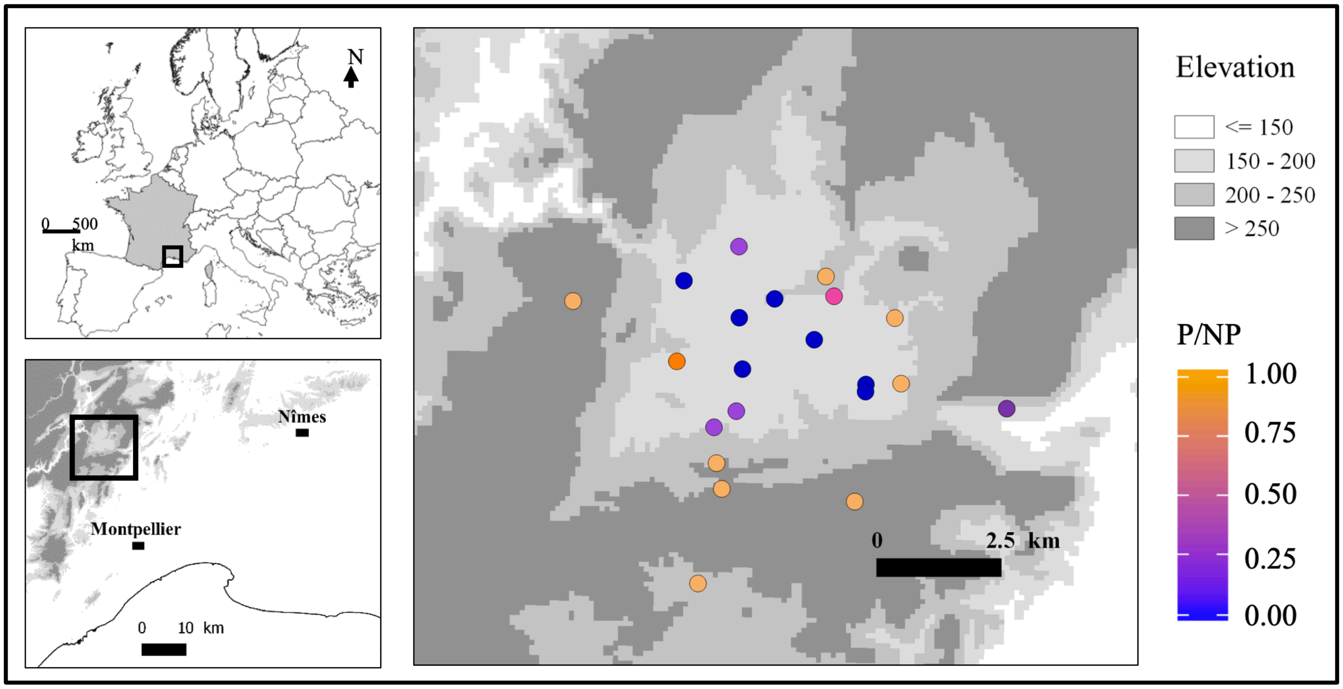


**Fig. S1. Map of study region and location of study sites.**

Color scale indicates the proportion of phenolic to non-phenolic individuals at each site.

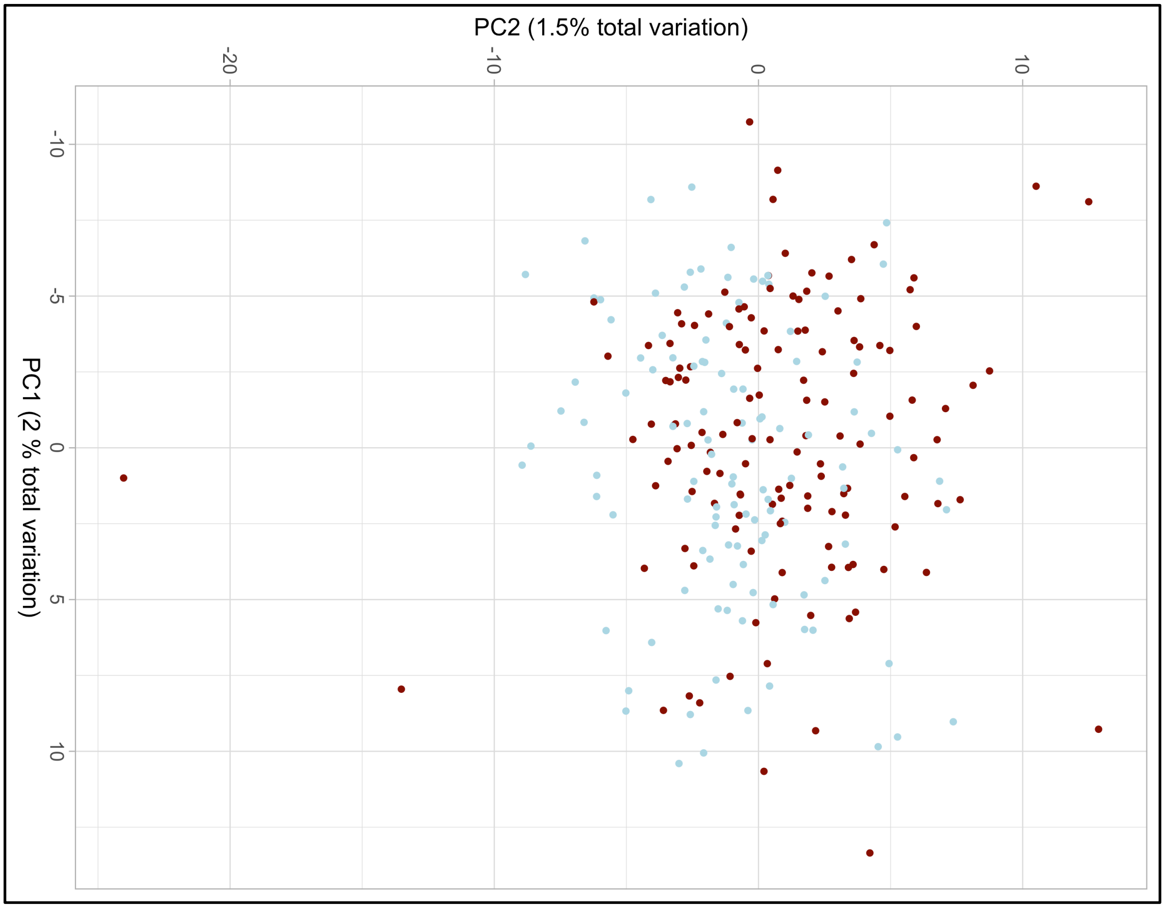


**Fig. S2. SNPs PCA and lack of genetic differentiation between ecotypes.**

Phenolic individuals (C or T chemotype) are colored in red. Non-phenolic individuals (G A U L chemotype) are colored in blue.

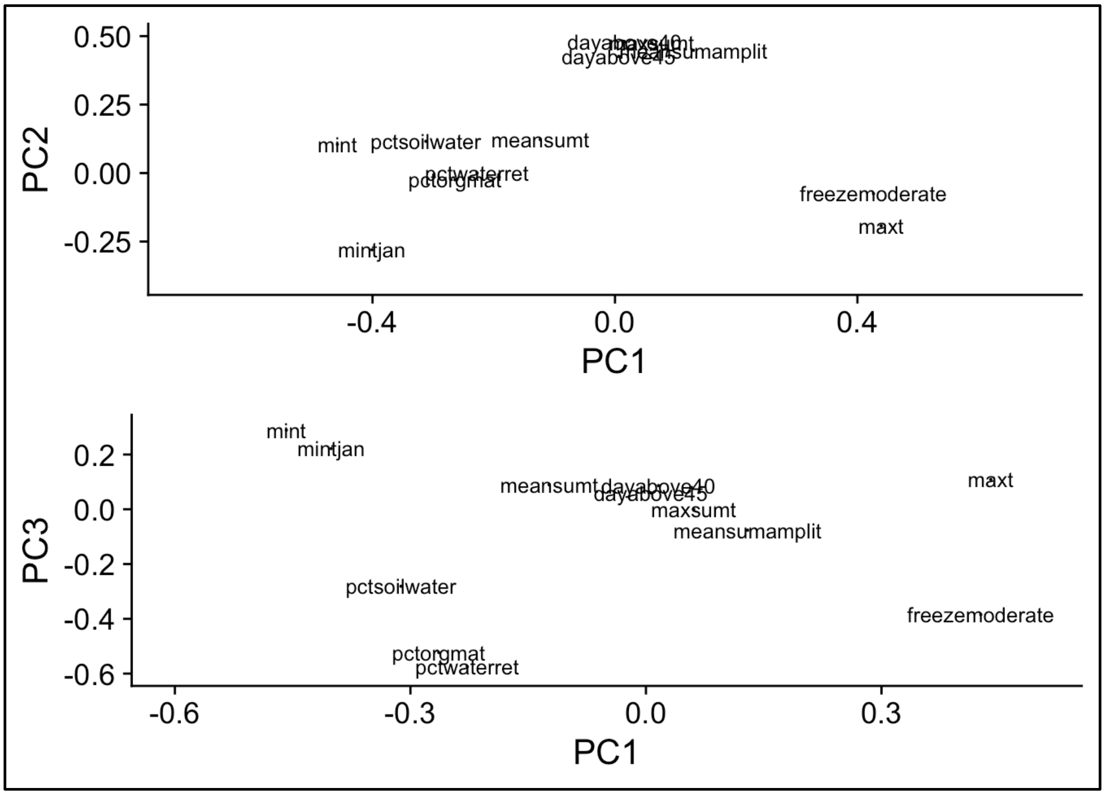


**Fig. S3. Projection of individual environmental variables on principal components of environmental variation.**

PC1, 2 and 3 account for 45%, 28% and 12% of the total environmental variation measured across sites. The individual variables used (centered and reduced to unit variance) were percentage water in soil, soil water retention and percentage organic matter in the soil ("pctsoilwater" "pctwaterret" "pctorgmat"), the mean daily minimum and maximum temperature in winter ("mint" "maxt"), the minimum temperature in the coldest month ("mintjan"), the mean daily minimum and maximum temperature the summer ("maxsumt" "meansumt"), the number of days where moderate to strong freezing was recorded (below -8C) "freezemoderate", the number of days exceeding 40C or 45C ("dayabove45" "dayabove40"), and the mean summer daily temperature amplitude ("meansumamplit"). Note that several of these individual variables are highly correlated as illustrated by their overlap in PCs plots.

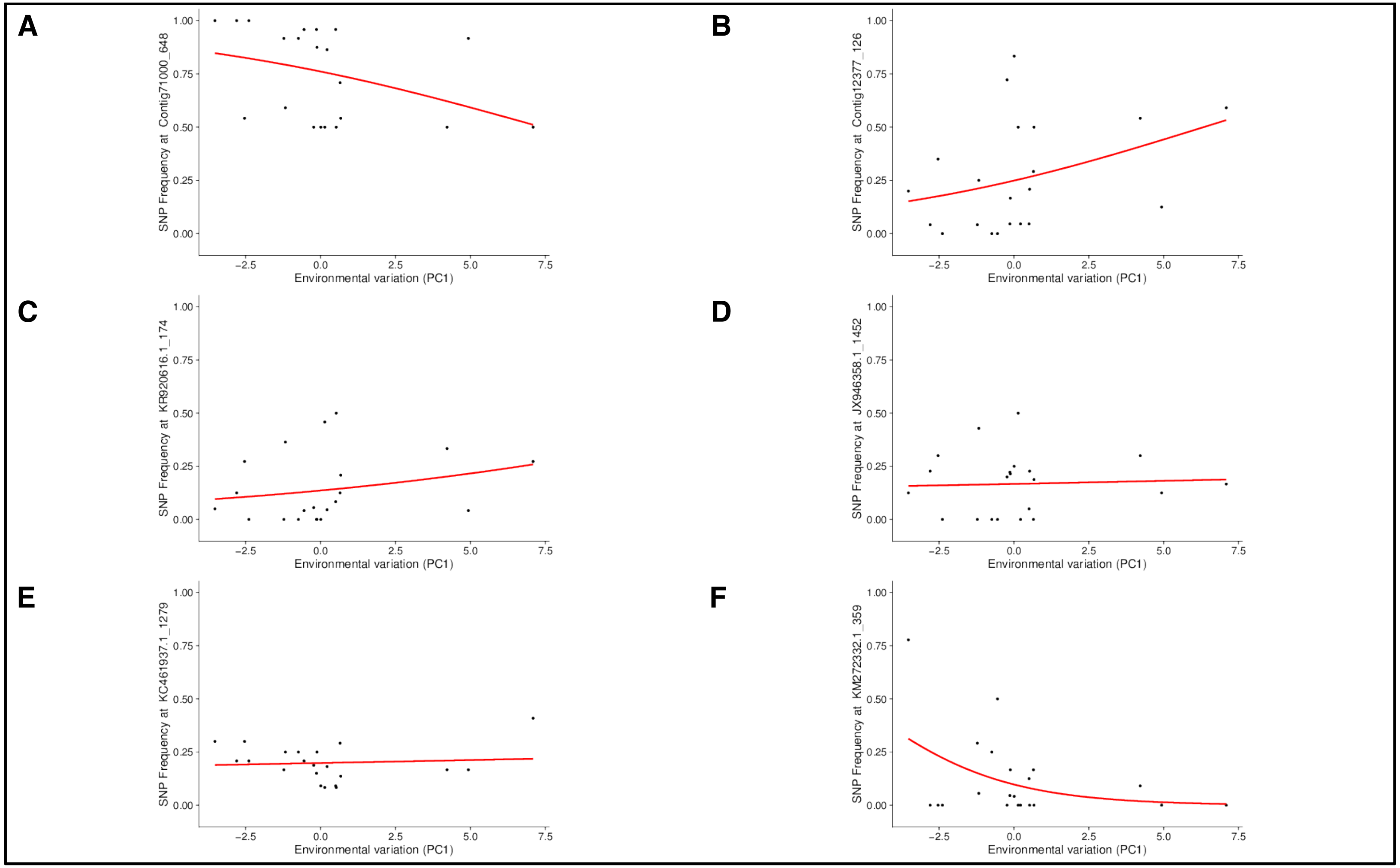


**Fig. S4.**

SNPs frequency clines along environment gradient at 6 SNPs explaining ecotype and chemotype identity. Environmental variation is measured using the first principal component of environmental variation. See supplementary Table 4 for R^2 and p-values associated with each model.

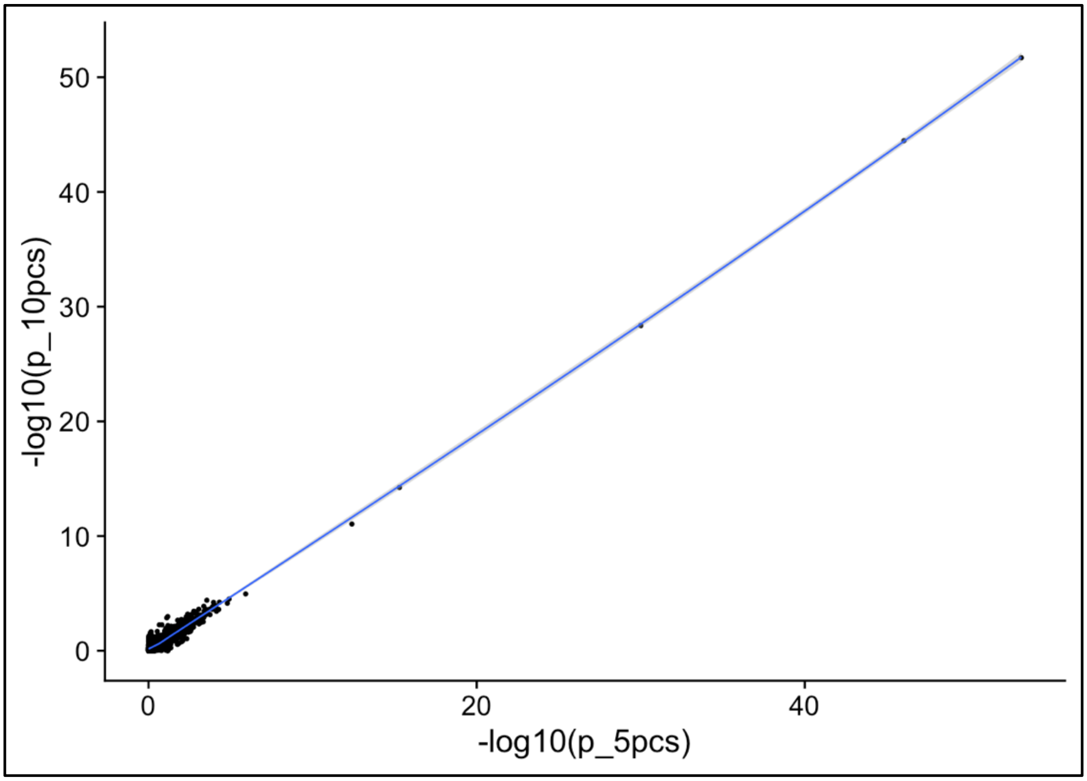


**Fig. S5. Robustness of PCs correction for population structure when detecting associations.**

We plot the correlation between -log10(p-values) for ecotype-SNP associations when using either 5 (p_5pcs) or 10 PCs (p_10PCs) of SNP variation to correct for background population structure. Blue lines indicate a “loess” non-parametric regression. Each dot indicates an individual SNP.

**Table S1. Repeatability of SNPs genotyping**

| Principal Component | Repeatability[$] | Cumulative $R^2$ |
|---|---|---|
| PC1 | 0.998 | 0.020 |
| PC2 | 0.999 | 0.035 |
| PC3 | 0.996 | 0.049 |
| PC4 | 0.995 | 0.062 |
| PC5 | 0.999 | 0.076 |
| PC6 | 0.998 | 0.088 |
| PC7 | 0.995 | 0.100 |
| PC8 | 0.996 | 0.111 |
| PC9 | 0.990 | 0.123 |
| PC10 | 0.992 | 0.134 |

$ Repeatability is calculated by Pearson's product-moment correlation coefficient of each PC coordinate using the coordinates of 23 independent paired technical replicates.

Cumulative $R^2$ indicates the proportion of the total variance explained by PC1, PC1+ PC2, etc.

**Table S2. Genotypes at the top 6 candidate loci**

| Gene / snp | n | ref | alt | freq_alt | nb0 | nb0_HW | nb1 | nb1_HW | nb2 | nb2_HW | Fis | p_HW |
|---|---|---|---|---|---|---|---|---|---|---|---|---|
| Contig71000_648 | 250 | T | C | 0.75 | 0 | 16 | 120 | 93 | 120 | 140 | -0.34 | 1.4e-07 |
| Contig12377_30 | 200 | A | T | 0.15 | 140 | 140 | 60 | 51 | 0 | 4.6 | -0.18 | 0.011 |
| KR920616.1_174 | 240 | A | T | 0.14 | 170 | 180 | 69 | 59 | 0 | 4.9 | -0.17 | 0.0099 |
| JX946358.1_1452 | 190 | T | C | 0.17 | 120 | 130 | 62 | 52 | 0 | 5.2 | -0.2 | 0.0064 |
| KC461937.1_1279 | 240 | G | A | 0.2 | 140 | 150 | 93 | 75 | 0 | 9.2 | -0.25 | 0.00016 |
| KM272332.1_359 | 240 | T | A | 0.11 | 200 | 190 | 28 | 48 | 13 | 3.1 | 0.41 | 1.7e-10 |

n: number of individuals genotyped

ref and alt denote the two SNP alleles.

nbi and nbi_HW, with i in {0,1,2}, denote respectively the observed and expected number of individuals in each genotype category i. Expected counts assume random mating and Hardy-Weinberg proportions

Fis denotes the deficit or excess of homozygotes relative to Hardy-Weinberg proportions and is computed as Fis =1 – n1/n1_HW.

p_HW is p_value of a goodness of fit test for observed versus Hardy-Weinberg expected proportions (note that these are not Bonferoni corrected).

**Table S3: Segregating redundancy (SR) of ecotypes and chemotypes**

| Ecotypes | SR | SE(SR) |
|---|---|---|
| Phenolic | 2.42 | 0.06 |
| Non-phenolic | 2.14 | 0.03 |
| **Chemotypes** | | |
| G | 2.28 | 0.08 |
| A | 3.60 | 0.52 |
| U | 1.55 | 0.11 |
| L | 3.78 | 0.79 |
| C | 7.23 | 0.97 |
| T | 3.67 | 0.31 |

Segregating redundancies are computed using the frequencies of the 3 and 6-loci genotypes and the genotypes to phenotypes relationships pictured in Fig 1 and 2.

**Table S4: Effect of environment on change in allele frequency at 6 SNPs with the strongest associations to ecotype and chemotype identity.**

| Gene / SNP position | Beta1 | R2 env1 | p1 | Beta2 | R2 env2 | p2 |
|---|---|---|---|---|---|---|
| Contig71000 / 648 | -1.475 | 0.182 | 0.138 | -0.123 | 0.001 | 0.902 |
| Contig12377 / 126 | 3.476 | 0.161 | 0.001 | 2.000 | 0.056 | 0.042 |
| KR920616.1 / 174 | 3.452 | 0.189 | 0.001 | -2.655 | 0.112 | 0.008 |
| JX946358.1 / 1452 | 0.875 | 0.018 | 0.386 | -1.845 | 0.078 | 0.068 |
| KC461937.1 / 1279 | 1.079 | 0.113 | 0.287 | 2.190 | 0.532 | 0.021 |
| KM272332.1 /359 | -1.412 | 0.040 | 0.142 | 2.143 | 0.098 | 0.022 |

Beta1 (respectively Beta2) is the standardized slope (Beta = slope /SE(slope) measuring the effect of environmental variation (respectively envPC1 and envPC2) on SNP allele frequency (as fitted by a logistic regression, see methods).

R2 env1 (respectively R2 env2) measures the portion of variance in allele frequency explained the model relative to a background model without the envPC1 (envPC2). R2 are obtained from ratio of deviance of logistic regression models (with and without the envPCs as predictor, see methods)

p1 (respectively p2) is the p-value for the likelihood ratio test of association of a focal SNP with envPC1 (respectively envPC2). P-values in bold are those <0.05/(2x6) = 0.00417.

**Dataset S1 (separate file).** Capture Adapters.xlsx.

Excel file containing the sequence of the specific hexamer barcodes used for genomic capture (see supplementary text below)

### Supplementary Information text for methods: locus targeted sequencing protocols

#### Plant DNA purification

DNA was extracted from 15 mg of fresh young leaves with the Chemagic DNA Plant Kit (Perkin Elmer Chemagen, Baesweller, DE, Part # CMG-194), according to the manufacturer's instructions. The protocol is adapted to the use of the KingFisher Flex™ (Thermo Fisher Scientific, Waltham, MA, USA) automated DNA purification workstation.

#### Construction of enriched library and sequencing

Genomic library preparation for multiplexed individuals and enrichment step by capture follow published protocols (1,2) with some modifications. The baits where designed bioinformatically to target a set of candidate genes identified by transcriptome sequencing (3) as well as candidate for known genes encoding enzymes of the monoterpene biosynthesis pathway previously identified in T vulgaris.

##### A Target preparation, construction of barcoded genomic libraries

1 : For each individual, 1 µg of total DNA (in 100 µL of water) are sheared using a Bioruptor Pico (Diagenode, Seraing, BE) sonication device in 500 µl microtubes to a targeted 300 bp DNA fragment size using parameters of the 300pb standard protocol for DNA shearing. *(*https://www.diagenode.com/files/protocols/Standard_protocols_for_DNAShearing.pdf*).*

2: 400 ng of fragmented DNA (in 40 µl of water) are blunted and 5' phosphorylated using the Thermo Scientific Fast DNA End Repair Kit (Thermo Fischer Scientific, Waltham, MA, USA, Part # K0771). A clean-up step is performed with 1 x volume of Agencourt AMPure XP magnetic beads. The elution volume is 20 µL.

3: Fragmented and repaired DNA are individually controlled (sizing and estimation of the concentration) by electrophoresis on a AATI Fragment Analyzer™ (Advanced Analytical Technologies, Ankeny, IA, USA) device with the DNF-474 High Sensitivity Fragment Analysis Kit.

4: 50 ng of fragmented DNA are ligated with 4 pmol of PE-P5 and MPE-P7 adapters. Each PE-P5 and each MPE-P7 adapter carries the same specific hexamer barcode (2), see file Capture Adapters.xls in Dataset S1) Reactions are conducted in 15 µl final volume with 1 unit of T4 DNA ligase for 1 hour at 22 °C followed by a heat inactivation step at 65°C for 10 minutes.

5: 48 samples (corresponding to 48 hexamer barcodes on the PE-P5 and PE-P7 adapter) are pooled. A clean-up step is performed with 1.8x volume of Agencourt AMPure XP magnetic beads. The elution volume is 94µL.

6: A nick fill-in step is performed using 64 units of Bst DNA polymerase (New England Biolabs, Ipswich, MA, USA, Part # M0275), 1x ThermoPol® reaction buffer, 250 µM dNTP in 120 µl final volume and incubated for 15 minutes at 37°C. A clean-up step is performed with 1.8x volume of Agencourt AMPure XP magnetic beads. The elution volume is 40 µL.

7: For each pool of 48 samples, a pre-hybridization PCR is performed using the Phusion® High-Fidelity PCR Master Mix (Thermo Fischer Scientific, Part # 1040-2678) with 200 nM PreHyb-PE_F (CTTTCCCTACACGACGCTCTTC) and 200 nM PreHyb-MPE_R (TGACTGGAGTTCAGACGTGTG) primers in a final volume of 100µl.

Thermocycling parameters: 3 minutes at 98°C, followed by 12 cycles of 80 seconds at 98°C; 45 seconds at 55°C and 60 seconds at 68°C, with a final elongation of 10 minutes at 72°C. A clean-up step is performed with 1.8x volume Agencourt AMPure XP magnetic beads. The elution volume is 20 µL.

#### B Enrichment, capture by hybridisation

The protocol used is based on Mascher et al (2), the User Manual of the MYBaits Sequence Enrichment for Targeted Sequencing kit (http://www.mycroarray.com/pdf/MYbaits-manual-v2.pdf) and on the Roche NimbleGen SeqCap EZ Library SR User's Guide (http://sequencing.roche.com/products/nimblegen-seqcap-target-enrichment/seqcap-reagents.html)

##### B-1 First round of hybridization of the barcoded libraries to biotinylated RNA probes

8: Prior to hybridization, 10 µl of Roche Diagnostics (Indianapolis, IN, USA) proprietary SeqCap EZ Developer Reagent (Roche, Part # 06684335001) were added to a 1.5-ml tube containing 0,5 µg of the 48 barcoded samples genomic library.

Next were added as blocking oligos:

- 1 µl (100 pmol/µl solution) of the P5 adapter blocking oligo , 5'-AGATCGGAAGAGCGTCGTGTAGGGAAAG
- and 1 µl (100 pmol/µl solution) of the MP7 adapter blocking oligo, 5'-AGATCGGAAGAGCACACGTCTGAACTCCAGTCA,

designed to block the truncated segment of TruSeq DNA library adapters during the sequence capture.

The mixture was dried down in a SpeedVac at 43°C during 20 to 30 min.

9: 7.5 µl of 2 × Sequence Capture Hybridization Buffer (tube 5, SeqCap EZ Hybridization and Wash Kit, Roche, Part # 05634261001) and 3 µl of Hybridization Component A (tube 6, SeqCap EZ Hybridization and Wash Kit, Roche, Part # 05634261001) were added. The hybridization cocktail was vortexed for 10 sec and collected by centrifugation. Following denaturation in a heat block (95°C, 10 min) the sample was transferred to a 0.2 ml PCR tube containing 80 ng of biotinylated RNA probes (4.5 µL of MYBaits Capture Probe).

10: The hybridization sample (15 µl) was incubated in a thermocycler (lid heated to 57°C) at 47°C for 64 h.

#### B-2 First round of washing of the captured library.

11: Streptavidin coupled magnetic beads are previously equilibrate as recommended by the Roche-Nimblegen protocol. Invitrogen Dynabeads MyOne™ Streptavidin C1 (Invitrogen, Thermo Fischer Scientific, Part # 65001) at 10 µg/µl were thoroughly vortexed, aliquoted (50 µl per hybridization) into 1.5-ml tubes and prepared for the affinity purification of captured DNA. The tubes were placed in a DynaMag-2 magnet (Invitrogen, Part # 123-21D) for 2 min. The clear liquid was discarded and 100 µl of 1 X Bead Wash Buffer (Tube 7, SeqCap EZ Hybridization and Wash Kit, Roche, Part # 05634261001) were added. Tubes were vortexed, placed back in the magnet, the clear liquid was removed, and the washing was repeated once. Dynabeads were resuspended in 50 µl 1 x Bead Wash Buffer, transferred into PCR plates and collected using a Agencourt SPRIPlate 96R (Agencourt, Part # A32782) . The clear supernatant was discarded.

12: The hybridization sample was added to the wet Dynabeads and mixed thoroughly by pipetting up and down. Using a thermocycler (lid heated to 57°C) at 47°C for 45 min the captured sample was bound to the Dynabeads. The sample was vortexed for 3 sec in 15-min intervals to ensure that the Dynabeads remain in suspension. Dynabeads plus bound DNA (15 µl) were washed by adding 100 µl 1 X Wash Buffer 1 (pre-heated to 47°C for 1 h) (Tube 1, SeqCap EZ Hybridization and Wash Kit, Roche, Part # 05634261001) and vortexing for 10 sec.

13: The suspension was transferred to a 1.5-ml tube and placed in a DynaMag-2 device, and the supernatant was discarded once clear. Washing was continued by adding 200 µl 1 X Stringent Wash Buffer (pre-heated to 47°C for 1 h) (Tube 4, SeqCap EZ Hybridization and Wash Kit, Roche, Part # 05634261001) The sample was mixed by pipetting avoiding a major temperature drop and incubated for 5 min at 47°C. The tube was placed in the DynaMag-2 magnet, the liquid was discarded and the washing at 47°C with 1 X Stringent Wash Buffer was repeated once.

14: 200 µl 1 X Wash Buffer 1 (pre-heated to room temperature) was added to the Dynabeads plus bound DNA. The sample was vortexed for 2 min and the liquid was collected to the tube's bottom. Following magnetic concentration the liquid was discarded, and the sample was washed at room temperature with 200 µl 1 X Wash Buffer 2 (vortexing for 1 min) (Tube 2, SeqCap EZ Hybridization and Wash Kit, Roche, Part # 05634261001), followed by a wash with 200 µl 1 X Wash Buffer 3 (vortexing for 30 sec) (Tube 3, SeqCap EZ Hybridization and Wash Kit, Roche, Part # 05634261001) as described for washing with Wash Buffer 1. The tube was removed from the magnet, the bead-bound captured library was resuspended in 25 µl PCR-grade water and the entire sample (beads + liquid) was transferred to a 0.2 ml PCR tube.

**B-3 Second round of hybridization of the barcoded libraries to biotinylated RNA probes**

15: The captured sample (25 µl) was denatured by incubation in a thermocycler (lid heated to 105°C) at 95°C for 3 min. 21 µl of the denatured solution were quickly transferred on to a 1,5 ml microtube.

16: Next were added to the captured sample

- 1 µl (10 pmol/µl solution) of the P5 adapter blocking oligo
- 1 µl (10 pmol/µl solution) of the MP7 adapter blocking oligo,
- 1 µl of SeqCap EZ Developer Reagent.

The mixture was dried down in a SpeedVac at 43°C during 10 to 20 min.

17: 7.5 µl of 2 × Sequence Capture Hybridization Buffer (tube 5, SeqCap EZ Hybridization and Wash Kit) and 3 µl of Hybridization Component A (tube 6, SeqCap EZ Hybridization and Wash Kit) were added. The hybridization cocktail was vortexed for 10 sec and collected by centrifugation. Following denaturation in a heat block (95°C, 10 min) the sample was transferred to a 0.2 ml PCR tube containing 15 ng of biotinylated RNA probes (1 µL of MYBaits Capture Probe) and 3,5 µl of UP water.

18: The hybridization sample (15 µl) was incubated in a thermocycler (lid heated to 57°C) at 47°C for 20 h.

**B-2 Second round of washing of the captured library.**

19: Streptavidin coupled magnetic beads are prepared as previously described (#11)

20: The hybridization sample was added to the wet Dynabeads and mixed thoroughly by pipetting up and down. Using a thermocycler (lid heated to 57°C) at 47°C for 45 min the captured sample was bound to the Dynabeads. The sample was vortexed for 3 sec in 15-min intervals to ensure that the Dynabeads remain in suspension. Dynabeads plus bound DNA (15 µl) were washed by adding 100 µl 1 X Wash Buffer 1 (pre-heated to 47°C for 1 h) (Tube 1, SeqCap EZ Hybridization and Wash Kit) and vortexing for 10 sec.

21: The suspension was transferred to a 1.5-ml tube and placed in a DynaMag-2 device, and the supernatant was discarded once clear. Washing was continued by adding 200 µl 1 X Stringent Wash Buffer (pre-heated to 47°C for 1 h) (Tube 4, SeqCap EZ Hybridization and Wash Kit) The sample was mixed by pipetting avoiding a major temperature drop and incubated for 5 min at 47°C. The tube was placed in the DynaMag-2 magnet, the liquid was discarded and the washing at 47°C with 1 X Stringent Wash Buffer was repeated once.

22: 200 µl 1 X Wash Buffer 1 (pre-heated to room temperature) was added to the Dynabeads plus bound DNA. The sample was vortexed for 2 min and the liquid was collected to the tube's bottom. Following magnetic concentration the liquid was discarded, and the sample was washed at room temperature with 200 µl 1 X Wash Buffer 2 (vortexing for 1 min) (Tube 2, SeqCap EZ Hybridization and Wash Kit), followed by a wash with 200 µl 1 X Wash Buffer 3 (vortexing for 30 sec) (Tube 3, SeqCap EZ Hybridization and Wash Kit) as described for washing with Wash Buffer 1. The tube was removed from the magnet, the bead-bound captured library was resuspended in 22 µl PCR-grade water and the entire sample (beads + liquid) was transferred to a 0.2 ml PCR tube.

**C PCR post-capture and sequencing**

23: An on- beads PCR amplification is undertaken to enrich library fragments, extend the adaptor sequence and incorporate an index to the P7 adaptor. The PCR reaction is using the KAPA® HiFi HotStart ReadyMix PCR Kit (KAPABiosystems, Boston, MA, Part # KR0370) in à final volume of 50 µl with:
15 pmol of SOL-PE-PCR_F primer (1)
AATGATACGGCGACCACCGAGATCTACACTCTTTCCCTACACGACGCTCTTC
15 pmol of SOL-MPE-INDX_R indexed primers
CAAGCAGAAGACGGCATACGAGATXXXXXXGTGACTGGAGTTCAGACGTGT
This primer carries 6 bases of the official TruSeq Illumina Index.

Thermocycling parameters: 2 minutes at 98°C, followed by 18 cycles of 20 seconds at 98°C; 30 seconds at 62°C and 30 seconds at 72°C, with a final elongation of 5 minutes at 72°C. The reaction volume is 50 µL. A clean-up step is performed with 1.8x volume Agencourt AMPure XP magnetic beads. The elution volume is 20 µL.

24: Indexed libraries are individually controlled (sizing and estimation of the concentration) by electrophoresis on an AATI Fragment Analyzer™ device with the DNF-474 High Sensitivity Fragment Analysis Kit.

25: Three indexed libraries, corresponding to 144 captured barcoded DNA samples, are equally mixed. The final pooled library is quantified by qPCR with the KAPA Library Quantification Kit (Part # KK4824) and provided to the Get-PlaGe core facility (GenoToul platform, INRA Toulouse, France http://www.genotoul.fr) for sequencing.

26: The final pooled library is sequenced using the Illumina paired-end protocol on a single lane of a HiSeq3000 sequencer, for 2 x 150 cycles.
